# Functional connectivity in northern swamp deer (*Rucervus duvaucelii duvaucelii*) population across a fragmented, human-dominated landscape along Gangetic Plains of north India: Implications for conservation in non-protected areas

**DOI:** 10.1101/2023.04.05.535741

**Authors:** Shrutarshi Paul, Sohini Saha, Parag Nigam, Garima Pandey, Bilal Habib, Dhananjai Mohan, Bivash Pandav, Samrat Mondol

## Abstract

The Indian subcontinent has witnessed disproportionate declines in large mammalian herbivore communities. The northern swamp deer (*Rucervus duvaucelii duvaucelii*) exemplifies the conservation challenges of typical non-protected area species, where apart from distribution status other ecological information is limited for the upper Gangetic basin population. We combined elements of radio-telemetry and conservation genetics to evaluate dispersal patterns, population connectivity and assess genetic variation and inbreeding status of this population living across a highly human-dominated area. We genetically identified 266 unique swamp deer and further analyses revealed presence of two spatially-admixed genetic lineages with moderate heterozygosity (Ho=0.51, SD= 0.10) and low inbreeding (FIS=0.133) status. Multi- disciplinary evidence suggests that the small, isolated grassland patches between Jhilmil Jheel Conservation Reserve (JJCR) and Hastinapur Wildlife Sanctuary (HWLS) are highly preferred by swamp deer during migrations and are genetically connected. The southern part of the area in HWLS showed early signatures of genetic discontinuity that require immediate conservation attention. We hypothesized that the human settlement history of this landscape, river dynamics and species’ ability to negotiate various pressures and disperse has helped to maintain such connectivity. While these signatures are encouraging for this small, isolated cervid population, careful management interventions are required to ensure the integrity and functionality of this landscape. We recommend a scientifically robust population estimation approach across this landscape and a multi-stakeholder-driven strategies to augment population and habitat recovery, plantation and riverscape management to ensure long-term survival of this species.

## 1. Introduction

The wild large mammalian communities have experienced severe population declines during last century (Ceballos et al. 2010; Ripple et al. 2015). Largely driven by various anthropogenic factors including overexploitation of natural resources, habitat loss and hunting (Isaac et al. 2007; Morrison et al. 2007), large proportions (∼50%) show reduced population size whereas ∼25% are threatened with extinction (Schipper et al. 2008; Karanth et al. 2010). The conservation challenges are further exacerbated due to complex and inherent life-history strategies, habitat-specificity, endemism and outside protected area distributions in many species (Harihar 2011; Ripple et al. 2016; Punjabi and Rao 2017; Dorji et al. 2019). In particular, survival of the species residing within human-dominated landscapes will depend on critical assessment of important factors that govern their ecology, demography and other biological parameters across their distributions. The Indian subcontinent, being considered as mega-biodiversity hotspot retains a large number of such conservation-concerned species. Particularly the large herbivore assemblages are facing strong impacts of anthropogenic pressures where ∼80% of the species are classified as ‘Threatened’ by IUCN (Ripple et al. 2015). The habitat-specialist herbivores (for example, rhinoceros, wild buffalo, swamp deer, brow-antlered deer etc.) were designated as the most affected (Karanth et al. 2010) with recommendations for generating detailed information for their long-term conservation.

The obligate, grassland-dwelling swamp deer (*Rucervus duvaucelii*) is currently considered one of the most extinction-prone megaherbivore in the Indian subcontinent (Karanth et al. 2010). Once distributed across the riverine floodplains between Pakistan and Bangladesh through India, it is now restricted to isolated pockets in some parts of north, north-east and central India and south-west Nepal (Schaller 1967; Groves 1982; Sankaran 1989; Qureshi et al. 2004). Out of the three known subspecies (Pocock 1943; Ellerman and Morridon-Scott 1951; Groves 1982; Qureshi et al. 2004), the northern counterpart (*Rucervus duvaucelii duvaucelii*) makes up ∼80% the global species population, distributed across two different areas in India namely the Sharda and Ganges habitat blocks spread through the northern states of Uttarakhand and Uttar Pradesh (Paul et al. 2020). The Sharda habitat block is part of the Terai Arc landscape whereas the Ganges habitat block represents the western-most distribution of the species (Qureshi et al. 2004; Paul et al. 2020). The swamp deer populations of the Sharda habitat block are relatively well studied and received adequate conservation attention as majority of them are found within protected areas (Qureshi et al. 2004; Ahmed and Khan 2008). On the other hand, the information on swamp deer from the Ganges habitat block has been insufficient till very recent time. Information on certain aspects of swamp deer ecology (group composition, feeding habits, activity budget- Tewari and Rawat 2013a, b, c, d, e; habitat assessment- Khan et al. 2003) was available from protected areas of the Gangetic habitat block. Recently, Paul et al. (2018, 2020) mapped the grassland habitats, documented detailed distribution and reported potentially small, inbred and scattered populations of the northern swamp deer across multiple fragmented grasslands covering both protected and non-protected areas in the upper part of the Gangetic habitat block. While this information has been critical in designating ‘Priority Conservation Areas’ in this habitat block (Paul et al. 2020), appropriate conservation planning would require further detailed assessments of migration routes, inbreeding status and genetic variation of this population. This is important as the Gangetic block swamp deer exhibit seasonal migratory behaviour (Paul et al. 2021) and no information on the genetic status and connectivity is available for this landscape so far.

In this paper, we investigate movement patterns and genetic status of the upper Gangetic population of the northern swamp deer. More specifically, our objectives were (1) to assess the genetic variation and inbreeding status of Gangetic swamp deer population and (2) evaluate their movement patterns and population connectivity using different approaches. We used a multidisciplinary approach through ecological surveys, radio-telemetry and genetic tools to address these questions. Our findings have critical conservation implications for this species and the grassland habitats within this human-dominated landscape.

## 2. Materials and Methods

### 2.1 Research permissions

All required permissions for fieldwork and sampling were accorded by the Forest Departments of Uttarakhand (Permit Nos: 1575/C-32 and 978/6-32/56) and Uttar Pradesh (Permit Nos.: 2233/23-2-12 and 3438/23-2-12). The radio-collaring permission was approved by the Ministry of Environment, Forest and Climate Change (MoEF&CC), Government of India and Uttarakhand Forest Department (Permit No: 1-76/2017WL). Ethical approvals were provided by the Uttarakhand Forest Department.

### 2.2 Study Area

This study was conducted in the upper Gangetic habitat block between Jhilmil Jheel Conservation Reserve (JJCR), Haridwar (in Uttarakhand) and southern boundary of Hastinapur Wildlife Sanctuary (HWLS), Uttar Pradesh (Fig. 1a). The area covers ∼120 km stretch of the Ganges river along with its tributaries Banganga and Solani. A maximum width of 8 km from both banks of all these rivers was considered as the survey regions as all swamp deer habitats were earlier reported within 5-6 km of the river banks (Paul et al. 2018, 2020). In total, the study area comprised ∼ 3173 km^2^ of habitat covering both protected (HWLS and JJCR- 1677 km^2^ area) as well as non-protected (1496 km^2^ area) regions. This landscape is one of the most densely populated areas of the entire country (population density of 1164 people/ km^2^ compared to national average of 382 people/ km^2^ (Cenus of India 2011)) with a mosaic of land use patterns including agricultural fields (76%), waterbodies (7%), forest (6%), grassland (6%), settlement (4%) and scrubland (1%) ( Paul et al. 2021). Despite such high human footprint, these habitats retain rich faunal biodiversity including swamp deer (*Rucervus duvaucelii duvaucelii)*, hog deer (*Axis porcinus*), nilgai (*Boselaphus tragocamelus*), fishing cat (*Prionailurus viverrinus*), Wild boar (*Sus scrofa*) and birds such as sarus crane (*Grus antigone*), black-necked stork (*Ephippiorhynchus asiaticus*), lesser adjutant (*Leptoptilos javanicus*), Pallas’s fish eagle (*Haliaeetus leucoryphus*) and bar-headed goose (*Anser indicus*) (Bashir et al. 2012; Grimmett et al. 2013).

**Fig. 1:**
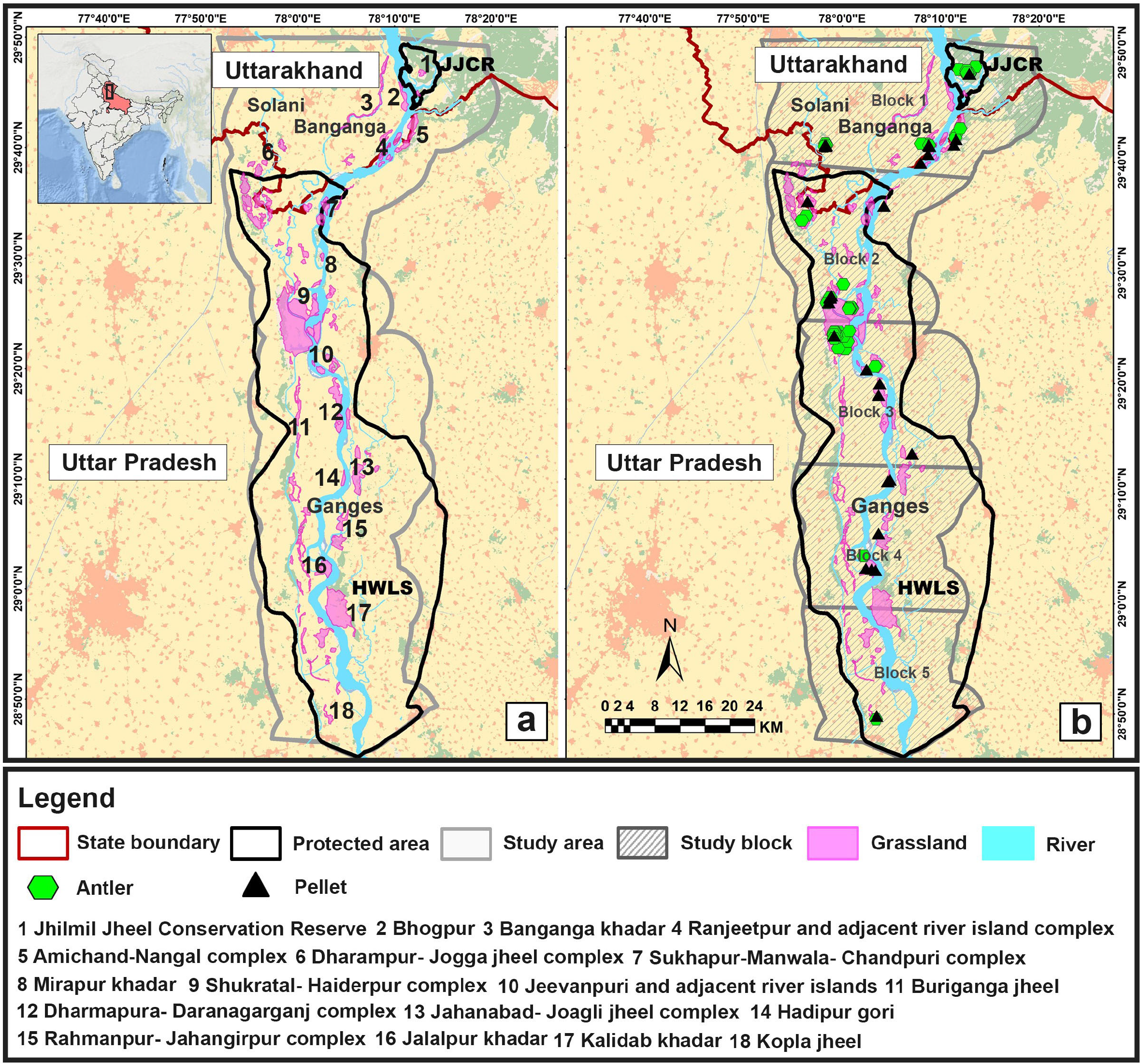
Representation of the study area and sampling efforts between Jhilmil Jheel Conservation Reserve (JJCR), Uttarakhand and Hastinapur Wildlife Sanctuary (HWLS), Uttar Pradesh. The section (a) (left panel) shows the protected areas (JJCR and HWLS) along with all the digitized grassland patches along the rivers Ganges and its tributaries Banganga and Solani. The section (b) (right panel) shows the locations of various types of samples used in this study within five distinct study blocks (24 km long stretches based on earlier recorded swamp deer movement patterns) along river Ganges.

### 2.3 Biological sampling, DNA extraction and species identification

We used earlier field-collected ungulate samples that were obtained as part of swamp deer surveys between JJCR and southern boundary of HWLS (Paul et al. 2018, 2020) in northern India. A total of 258 antlers and 2499 pellet samples were available to estimate genetic diversity of this population. We selected a total of 488 samples (258 antlers and 230 fresh pellets) for genetic analyses as they provided homogenous spatial representation of the study area ((Fig. 1b; Supplementary Table S1). We decided to use all the antler samples as they are known to provide good quality DNA for genetic analyses (Gupta et al. 2013; Venegas et al. 2020).

In the laboratory, we performed DNA extraction from the antler and pellet samples using already-established protocols (Paul et al. 2019). In brief, we cut each antler using individual sterile saw blades from the base and collected the powders in separate tubes. About 20 mg of the powder was weighed, decalcified (in 0.5M EDTA (pH 8) for 48 hours) followed by lysis (with 40μl Proteinase K and 400μl of ATL solution at 56°C for seven days). For pellets, we swabbed the top layer of each sample using sterile swabs (Biswas et al. 2019). In both cases, DNA extraction was done using spin-column protocol of the QIAamp DNA Tissue Kit (Qiagen Inc, Hilden, Germany) and DNA was eluted twice in 100μl preheated 1X TE buffer. Negative controls were kept to monitor any possible contamination. To reduce the contamination chances from poor-quality samples, all pellet extractions were performed in a physically separate laboratory space dedicated for non-invasive samples ensuring geographic separation between pre and post-PCR workspace. Antler DNA was extracted in a separate DNA extraction facility at the institute. In addition, standard procedures including regular sterilization (through UV and bleach) of the laboratory between processing of different batches of samples, inclusion of extraction and PCR negatives etc. were maintained.

The antler samples did not require any molecular species identification due to their distinctive morphological features (six-tined patterns) (Qureshi et al. 2004). However, we used swamp deer-specific molecular assays for pellet DNA (Paul et al. 2019) to remove other co-existing ungulates from further genetic analyses. The PCR reactions (10μl volume) contained 4μl multiplex buffer (QIAGEN Inc.), 4μg of BSA (4mg/ml), 0.25 μM of primer mix and 2μl each of 1:10 diluted DNA extracts and DNAse-RNAse free water with following conditions: initial denaturation (95°C for 15 min); 45 cycles of denaturation (95°C for 30 sec), annealing (50°C for 40 sec) and extension (72°C for 40 sec), followed by a final extension (72°C for 10 min). Negative controls were included to monitor contamination. The amplified products were checked in 2% agarose gel for species-specific band patterns (Paul et al. 2019). Samples not confirmed with swamp deer-specific markers were subjected to an ungulate-specific molecular assay (Gupta et al. 2014). Amplified products were cleaned with Exonuclease (Thermo Scientific, Waltham, USA) and Shrimp Alkaline Phosphatase (Amresco, Solon, USA) mixture and sequenced bidirectionally in an ABI 3500XL bioanalyzer (Applied Biosystems). The sequences were aligned, manually screened for any ambiguities and matched against the Genbank database.

### 2.4 Individual identification and molecular sexing

The swamp deer population along the upper Gangetic plains is known to be small, fragmented and potentially inbred (Paul et al. 2020), and thus it was important to develop a marker panel for individual identification with high statistical support. We selected a set of 48 microsatellite markers (developed for red deer, Coulson et al. 1998) based on available information on polymorphism, amplicon size and success in cross-species amplification. We adopted a cost-effective universal M13 primer-based approach (Schuelke 2000; Csencsics et al. 2010)) to screen these markers with a set of reference swamp deer samples (24 antlers and 18 genetically identified pellets). The shortlisting criteria included (i) amplification success, (ii) amplicon size, (iii) polymorphism, (iv) ease in allele calling and (v) stable allele characteristics (Hoffman and Amos 2005; Pompanon et al. 2005; Linacre et al. 2011; Johnson et al. 2014; Ghosh et al. 2021). PCR reactions were performed in 10μl volumes containing 4μl Qiagen multiplex PCR master mix (QIAGEN Inc.), 0.2μM forward primer, 0.1μM reverse primer, 0.2μM labelled M13 primer, BSA (4mg/ml) and 2μl of the DNA extract (1:10 diluted) with negative controls. PCR reactions included an initial denaturation at 95°C for 15 min; 45 cycles of denaturation at 95°C for 30 sec, annealing at 57°C for 40 sec, extension at 72°C for 40 sec; final extension at 72°C for 20 min. PCR products were genotyped (with HIDI formamide and Liz 500 size standard) in an automated sequencer ABI 3500XL (Applied Biosystems). To ensure good data quality each marker was amplified three independent times. Alleles were scored using program GENEMARKER (Soft genetics Inc., Pennsylvania, United States) and data quality was maintained by using ‘quality index’ approach (Miquel et al. 2006; Modi et al. 2018), where a quality score of 0.66 or above was approved. The final set of markers was individually labeled, standardised as multiplex reactions and compared (with the M13 data) for data consistency.

We used molecular sexing assay (Paul et al. 2019) to ascertain sex of the individually identified pellet samples. The PCR reactions contained 4μl multiplex buffer (QIAGEN Inc.), 4μg of BSA (4mg/ml), 0.25 μM of primer mix, 2μl each of 1:10 diluted DNA extracts and DNAse-RNAse free water. PCR conditions included an initial denaturation (95°C for 15 min); 45 cycles of denaturation (95°C for 30 sec), annealing (57°C for 40 sec) and extension (72°C for 40 sec), followed by a final extension (72°C for 10 min). Negative controls were included to monitor contamination. The amplified products were checked in 3% agarose gel for sex-specific band patterns (Paul et al. 2019).

### 2.5 Data analysis

#### 2.5.1 Genetic variation

We ascertained genetic recaptures by comparing all the genotype data in program CERVUS (Kalinowski et al. 2007). After removing the recaptures, we used program GIMLET (Valière 2002) to identify the low-frequency alleles (less than 10% in the entire data set) for confirmation and calculated PID(sibs) for the dataset. For the final microsatellite panel, we calculated locus-wise and overall summary statistics (GIMLET (Valière 2002) and genotyping error rates (MICROCHECKER v 2.2.369 (Van Oosterhout et al. 2004), FreeNA (Dempster et al. 1977; Chapuis and Estoup 2007) and data-based calculations (Broquet and Petit 2004)). Program ARLEQUIN (Excoffier et al. 2005) was used to determine Hardy-Weinberg equilibrium and linkage disequilibrium.

#### 2.5.2 Population structure detection

We used both Bayesian (STRUCTURE and GENELAND) as well as non-Bayesian (DAPC) approaches to infer patterns of swamp deer genetic structure within the study area. In STRUCTURE analyses, we performed 10 independent runs for each cluster value (K) between 1 and 10 with both non-spatial and locprior models, along with correlated allele frequency model. A total of 450,000 iterations and 50,000 burn-in were performed. For the locprior model-based analyses, we stratified the 120 km stretch of Ganges river in the study area into five continuous blocks (each block length ∼24 km) based on earlier recorded swamp deer movement patterns in this landscape (Paul et al. 2021). The optimal value of K was assessed using the “Evanno” method in STRUCTURE HARVESTER web version (Earl and vonHoldt 2012).The individuals were sorted as “pure” or “admixed” based on 70% ancestry coefficient threshold values (Ashrafzadeh et al. 2021). We also used another Bayesian clustering approach implemented in program GENELAND version 4.0.3 (Guillot et al. 2005) to assess spatial patterns of genetic structure without assuming admixture (Guillot et al. 2005; De et al. 2021). We used the spatial model assuming 10 clusters (with no uncertainty on coordinates), correlated allele frequencies and ran the analyses with 100000 iterations of which every 100^th^ observation was retained. Additionally, to test the extent of genetic structuring in this landscape, we used a non-Bayesian programme, Discriminant Analysis of Principal Components (DAPC) using R-package ‘adegenet’ (Jombart et al. 2010) in R studio 1.1.463. This approach transforms the genetic data into principal components, followed by clustering to define group of individuals with a consideration of minimum within-group variation and maximum between-group variations among the clusters. We used the apriori population cluster assignment approach (five locations as used in Structure locprior model) where we selected the number of Principal Components based on optimisation of ‘α’ score through spline interpolation (De et al. 2021). We calculated genetic differentiation (pairwise FST) in ARLEQUIN version 3.1 (Excoffier et al. 2005) between the genetic groups used in Structure analysis (Estes-Zumpf et al. 2010; He et al. 2010; Zachos et al. 2016).

#### 2.5.3 Inbreeding and relatedness analysis

We used individual-level genetic data to assess the degree of inbreeding in this population. We calculated inbreeding coefficient (FIS) using program FSTAT (version 2.9.3.2; (Goudet 2002), where p value was computed through 13000 randomisations (Gibbs et al. 1997; Heuertz et al. 2004; Hernández et al. 2020). Further, we used program ML-RELATE (Kalinowski et al. 2006) to calculate maximum likelihood estimates of pair-wise relatedness and relationship categories between individuals. We selected highly related individuals (relatedness value of >0.6) with 100% amplification at all loci and plotted their geographical distribution in this landscape to establish genetic signatures of movements (Rodzen et al. 2004; Puill-Stephan et al. 2009; D’Aloia et al. 2018).

### 2.6 Capture, collaring and telemetry analysis

We undertook radio-collaring study to understand movement patterns of swamp deer in this landscape. We collared two apparently healthy female swamp deer using drive net approach in JJCR, Uttarakhand during May-June 2018 (Kock et al. 1987; López-Olvera et al. 2009). The animals were acclimatised (for three months) to the conditions before collaring operations. Once captured in the net, the animals were blindfolded and administered with a mild dose of sedative Azaperone (40mg/ml dose) and fitted with GPS Vertex Plus satellite collars (Vectronic Aerospace). The collars were set to provide information on latitude, longitude, time and temperature at every 2-hour interval. The two collared animals were monitored for 14 months (Female 1) and 11 months (Female 2), respectively to understand their movement routes and identify the critical stopover sites (based on 10% of total locations in any site, (Sawyer and Kauffman 2011). We analysed various movement parameters (total distance travelled, longest distance from starting point, mean step length, mean speed etc.) in ArcGIS 10.2.2. using the ArcMET (version 3) (Wall et al. 2013; Wall 2014). We performed Net squared displacement (NSD) analysis to categorise the movement patterns into different classes (migratory, mixed migratory, nomadic, dispersal and home range) for each individual (Bunnefeld et al. 2011).

## 3. Results

Out of 230 field-collected faecal pellets, 159 were confirmed as swamp deer, making the total sample size as 417 (258 antlers and 159 pellet samples) for downstream analysis. The remaining samples either belonged to other herbivores (n=44, nilgai-22; hog deer-16; barking deer-3 and domestic goat-3) or did not produce any results (n=27, possibly due to poor DNA quality).

### 3.1 Microsatellite markers and swamp deer genetic diversity

During initial standardization, 27 of the initially selected 48 markers were rejected due to various reasons (no amplification- two loci, inconsistent amplification- five loci, multiple bands- 16 loci, non-specific bands- four loci). Further scrutiny of the remaining markers (n=21) revealed that two loci showed excessive stutter bands and low RFUs, three loci did not produce stable allele characteristics and three markers showed inconsistent results with antler and pellet DNA, resulting in a final microsatellite panel consisting 13 loci (Supplementary Table S2). During individual identification, samples with at least 10 loci data were considered based on a statistically strong PID(sibs) value of 1*10^-4^ (given that the global population of swamp deer is ∼5000 individuals, (Duckworth et al. 2015).

We generated 10 or more loci data from 298 samples (190 antlers and 108 pellet samples), which resulted in identifying 266 unique swamp deer individuals (from 168 antlers and 98 faecal pellets). Remaining 32 genotypes were identified as genetic recaptures (ranging from 1- 3 recaptures of already identified individuals). The standardized STR panel showed a mean 91.19% success rate (84.93-95.14% range) and low genotyping errors (mean frequency of null alleles, false alleles and mean allelic dropout rate was 0.06 (range 0-0.17), 0.09 (range 0.05- 0.15) and 0.09 (range 0.04-0.18), respectively). Overall, the panel was found to be moderately polymorphic with a mean of 6 alleles (SD 3.48, varying between 2-13 alleles) and expected and observed heterozygosity of 0.60 (SD 0.15) and 0.51 (SD 0.10), respectively. None of the loci deviated from Hardy-Weinberg equilibrium and we found no strong linkage disequilibrium. Loci-wise summary statistics are shown in Table 1. Molecular sexing of the faecal pellets (n=98) ascertained 47 female and 51 male samples (219 males and 47 females in total).

**Table 1:** Summary statistics of the 13 microsatellite loci used for population genetic analyses of swamp deer in this study.

| SL No. | Locus | Species | Success rate (%) | No of alleles | Allelic Size Range | H<sub>E</sub> | H<sub>O</sub> | Null allele | Allelic Dropout rate | False allele rate | P<sub>ID(sibs)</sub> |
| --- | --- | --- | --- | --- | --- | --- | --- | --- | --- | --- | --- |
| 1 | CEH53 | Swamp deer | 86.95 | 13 | 48 | 0.84 | 0.59 | 0.11 | 0.04 | 0.10 | 3.43*10<sup>-1</sup> |
| 2 | CEH33 | Swamp deer | 90.98 | 12 | 24 | 0.81 | 0.56 | 0.13 | 0.04 | 0.15 | 1.23*10<sup>-1</sup> |
| 3 | CEH82 | Swamp deer | 93.71 | 11 | 26 | 0.81 | 0.66 | 0.07 | 0.06 | 0.12 | 4.42*10<sup>-2</sup> |
| 4 | CEH56 | Swamp deer | 91.81 | 7 | 12 | 0.71 | 0.61 | 0.05 | 0.06 | 0.14 | 1.90*10<sup>-2</sup> |
| 5 | CEH35 | Swamp deer | 88.97 | 5 | 8 | 0.66 | 0.34 | 0.17 | 0.07 | 0.13 | 8.77*10<sup>-3</sup> |
| 6 | CEH50 | Swamp deer | 95.14 | 5 | 8 | 0.62 | 0.54 | 0.04 | 0.06 | 0.07 | 4.36*10<sup>-3</sup> |
| 7 | CEH52 | Swamp deer | 84.93 | 8 | 18 | 0.59 | 0.5 | 0.05 | 0.08 | 0.06 | 2.23*10<sup>-3</sup> |
| 8 | CEH58 | Swamp deer | 89.56 | 6 | 22 | 0.56 | 0.46 | 0.07 | 0.11 | 0.06 | 1.19*10<sup>-3</sup> |
| 9 | CEH75 | Swamp deer | 93.12 | 4 | 8 | 0.53 | 0.59 | 0 | 0.08 | 0.05 | 6.75*10<sup>-4</sup> |
| 10 | CEH71 | Swamp deer | 88.61 | 2 | 8 | 0.46 | 0.61 | 0 | 0.18 | 0.08 | 4.18*10<sup>-4</sup> |
| 11 | CEH47 | Swamp deer | 93.71 | 6 | 14 | 0.43 | 0.35 | 0.07 | 0.17 | 0.09 | 2.60*10<sup>-4</sup> |
| 12 | CEH43 | Swamp deer | 95.02 | 4 | 6 | 0.42 | 0.44 | 0 | 0.12 | 0.05 | 1.65*10<sup>-4</sup> |
| 13 | CEH51 | Swamp deer | 93.00 | 3 | 10 | 0.38 | 0.38 | 0.01 | 0.13 | 0.06 | 1.10*10<sup>-4</sup> |
| Mean |  |  | 91.19 | 6.61 |  | 0.6 | 0.51 | 0.06 | 0.09 | 0.09 |  |
| SD |  |  | 3.185 | 3.477 |  | 0.158 | 0.107 | 0.053 | 0.047 | 0.035 |  |

### 3.2 Population Structure

The STRUCTURE results (with locprior model) show three genetic lineages (based on ancestry admixture coefficient threshold of 70%) in the swamp deer population (K=3). These three genetic lineages can be divided into the following groups: Group I- 44 individuals, Group II-126 individuals and Group III- 16 individuals, respectively. In addition, remaining 80 individuals showed signatures of mixed genetic lineages (mostly between Groups I and II) (Fig. 2a). The non-spatial model indicated K=2, where 87, 81 and 98 individuals were assigned to Group I, Group II and Group III (admixed), respectively. The DAPC analysis (with five apriori groups) showed the first four groups were overlapping whereas the fifth group formed a distinct entity. When examined closely, we found that individuals forming Group I, II and the mixed group (from STURCTURE locprior analysis) and first four groups (from DAPC analysis) were found throughout the landscape. The Group III from STRUCTURE locprior and the fifth DAPC group was mostly restricted to southern part of HWLS below Bijnor Barrage (Blocks 4 and 5) (Figs. 2b, 2c). Taken together, we interpret that the swamp deer population between JJCR and HWLS is genetically connected. The genetic differentiation among these five groups ranged between 0.005-0.09 (Table 2). However, the results of the GENELAND analyses suggests two genetic clusters (mode of posterior distribution at K=2) across the study landscape: the first cluster corresponding with the STRUCTURE locprior results (Group I, II and the mixed lineages formed a single group) and the second cluster corroborated with the separate group (Group III in case of STRUCTURE locprior and fifth group as per the DAPC analysis) (Fig. 2d).

**Table 2:** Genetic differentiation (pairwise Fst) between five study blocks in the upper Gangetic plains, north India.

| **Genetic differentiation among five spatial blocks** |  |  |  |  |  |
| --- | --- | --- | --- | --- | --- |
| **Blocks** | **1** | **2** | **3** | **4** | **5** |
| **1** | 0 |  |  |  |  |
| **2** | 0.033* | 0 |  |  |  |
| **3** | 0.009* | 0.024* | 0 |  |  |
| **4** | 0.006* | 0.039* | 0.021* | 0 |  |
| **5** | 0.064* | 0.091* | 0.089* | 0.067* | 0 |

\*Significant at $P < 0.05$ using 10,000 randomizations

**Fig. 2:**
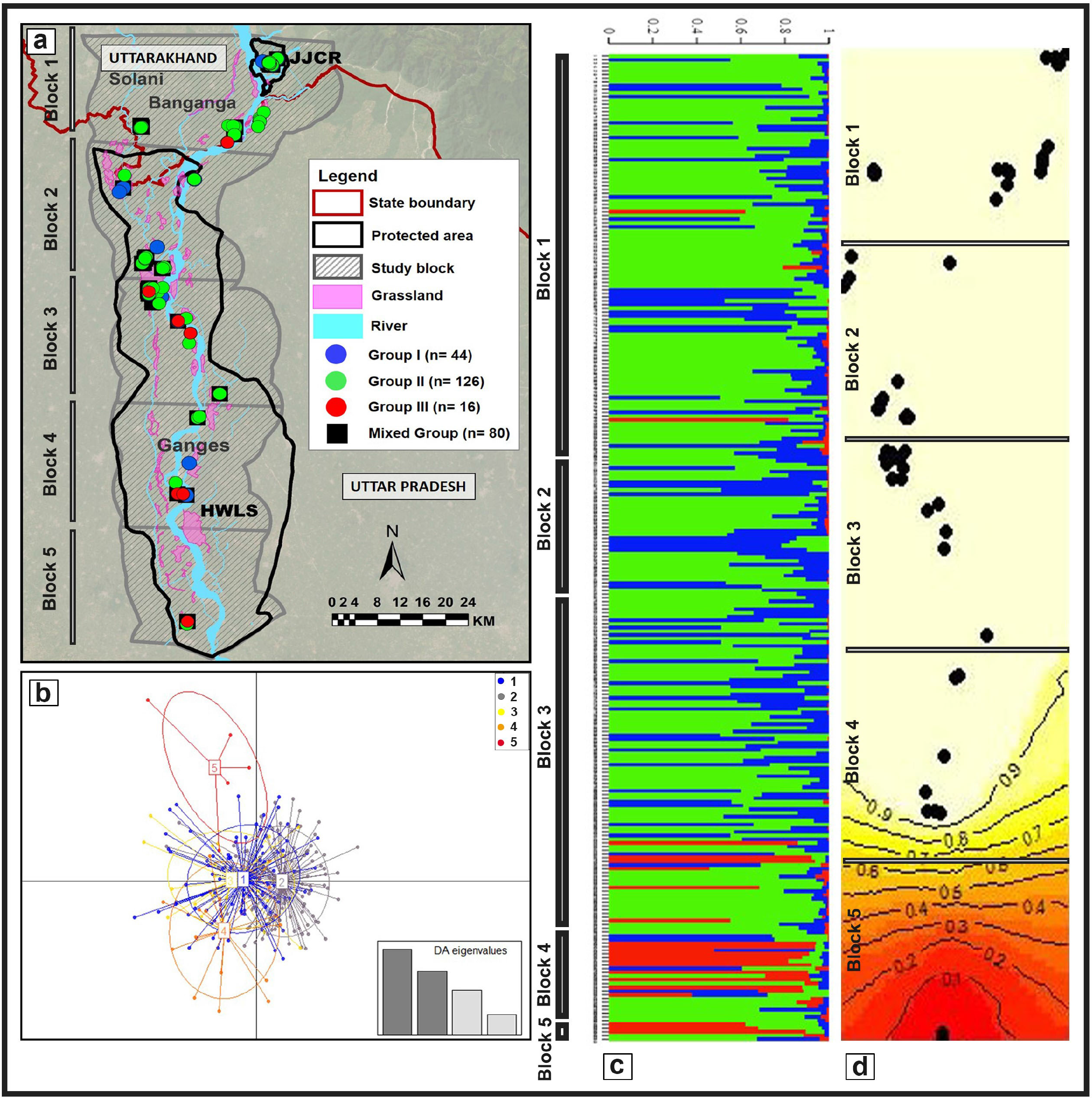
Outcomes of various Bayesian (STRUCTURE and GENELAND) and non-Bayesian (DAPC) genetic connectivity analyses for northern swamp deer. Panel (a) shows the distribution of different swamp deer genetic groups across this landscape; Panel (b) shows the DAPC results indicating genetic clusters (K=5) corresponding to the five study blocks; Panel (c) showing the genetic admixture patterns of the identified individuals based on sampled study blocks and; Panel (d) showing GENELAND outputs at K=2 where clear genetic discontinuity of study block 5 is depicted.

### 3.3 Inbreeding status

Based on the genetic data from the swamp deer individuals sampled in the upper Gangetic habitat (n=266), we found the inbreeding co-efficient (mean FIS value) to be 0.128 (p<0.05), indicating low levels of inbreeding. Overall, the males (n=219) show slightly higher FIS value of 0.175 (p<0.05) than the females (n=47, FIS- 0.133 (p<0.05)). Pair-wise average relatedness ranged between 0.0001-0.93 across the dataset. When the locations of all the highly-related individual pairs (relatedness>0.6, n=39 pairs) were plotted, eight pairs were found to be spread across the study landscape, supporting recent movement of individuals in this landscape (Supplementary Fig. S1).

### 3.4 Radio-collaring

Path trajectory analyses for both females (Female 1-14 months and Female 2- 11 months) showed a downward linear distance movement of 18 and 28 km for them, respectively. The maximum linear distance travelled, average step length, speed and other parameters for both the collared individuals are summarised in Table 3 (Figs. 3a, 3b). NSD analysis suggests that Female 1 exhibited mixed migratory (returning to a location situated midway between initial point and furthest point) whereas Female 2 showed migratory (returning back to its original location after 9 months) type movement patterns (Figs. 3c, 3d). Some of the important stopover points during swamp deer movement routes between JJCR and HWLS are grassland patches within Ranjeetpur and adjacent river island complex (Point c in Figs. 3a, 3b), Amichand- Nangal complex (Point d in Figs. 3a, 3b) and Sukhapur-Manwala-Chandpuri complex (Point e in Fig. 3b).

**Fig. 3:**
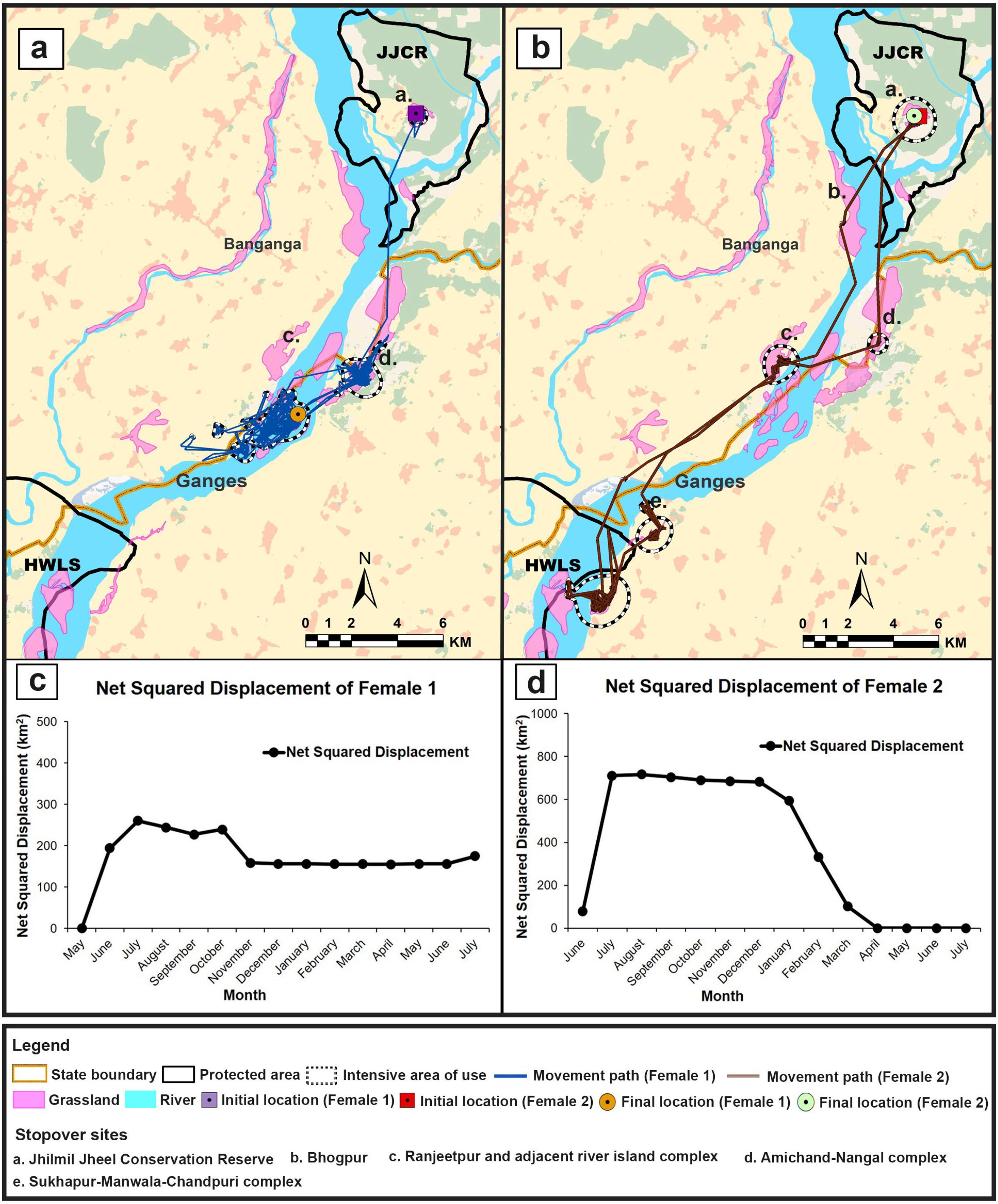
Movement patterns of two collared swamp deer females. Both panels (a) and (b) show the movement trajectory paths, intensive use areas and stopover sites along the river Ganges. Panels (c) and (d) shows the NSDs of the respective individuals depicting their migration types. **Supplementary Fig. 1:** Representation of spatial heterogeneity among some related individuals (r>0.6) sampled in our survey, indicating genetic connectivity and recent movement events within this landscape. Each pair of related individuals (total eight pairs) is indicated by same symbols presented in the figure.

**Table 3:** Basic movement parameters calculated for the two collared swamp deer females in this study.

| **Movement Parameters** | **Female 1** | **Female 2** |
| --- | --- | --- |
| Tracking duration (month) | 14 | 11 |
| Mean step length (km) | 0.117 | 0.082 |
| Mean speed (km/hr) | 0.058 | 0.040 |
| Maximum linear distance travelled from starting location (km) | 18 | 28 |
| Total distance (km) | 584.3 | 309.3 |
| Total displacement (km) | 15.2 | 0.227 |
| Migration type | Mixed migratory | Migratory |

## 4. Discussion

The densely-populated upper Gangetic plains of north India currently retain the westernmost distribution of swamp deer population that faces acute anthropogenic pressures in the form of habitat loss from rapid urbanisation, expanding human population and associated agricultural activities (Paul et al. 2020). As significant portion of these available habitats fall outside protected area jurisdictions, habitat conservation and population management is experiencing serious challenges. Further, scattered and inadequate information on their habitat use and movement patterns across the landscape make any conservation plan difficult. This study has generated probably the most exhaustive primary information on swamp deer dispersal patterns and genetic status in this landscape showcasing the importance of the remaining patchy grassland habitats for their future survival. Our assessments based on a combination of genetic and radio-telemetry approaches revealed that the entire fragmented landscape between JJCR and HWLS is functionally connected and the small, isolated patches of the grasslands are regularly used by swamp deer for their seasonal dispersals. These results substantiate the earlier research focused on the importance of the remnant grassland habitats from this landscape (Khan and Khan 1999; Khan et al. 2003; Qureshi et al. 2004; Tewari and Rawat 2013b; Paul et al. 2018, 2020; Mondol et al. 2019). Such combined approaches have been adopted in other studies to derive conclusions about connection between geneflow and movement patterns (Riley et al. 2006; Boulet et al. 2007; Kaczensky et al. 2011; Gustafson et al. 2017; Carvalho et al. 2018). One of the most important outcome of this study is identification of 266 northern swamp deer individuals within the Gangetic habitat block region. This is probably the first report of confirmed minimum numbers of this subspecies from this landscape. The latest assessment of the northern subspecies population size is reported as ∼3500 (Prakash et al. 2012; Duckworth et al. 2015; Wildlife Institute of India 2017; Islam et al. 2022), but the methods through which this assessment have been made (for example, direct count by foot and elephant back- Sinha et al. 2007; direct count elephant back and vehicle sampling- Ahmed and Khan 2008; focal sampling- Rastogi et al. 2023 method used in Jhilmil Jheel Conservation Reserve and Dudhwa Tiger Reserve) require detailed validations. There is an urgent need to compare different population estimation approaches for swamp deer and establish a reliable method for this. Results from such accurate population estimation across their distribution will strongly help in re-evaluating the species conservation status (currently considered as ‘Vulnerable’ by IUCN (Duckworth et al. 2015)) and help in their conservation. This is particularly important as significant portion of the northern swamp deer distribution is outside protected area regime, where conventional management/ conservation efforts based on Government regulations are ineffective. In this regard, future efforts should consider using standard genetic (microsatellites as in this study and others-Coulon et al. 2006 Frantz et al. 2006Miotto et al. 2011Atterby et al. 2015 Vergara et al. 2015) or genomic (SNPs-Edea et al. 2013; Viengkone et al. 2016; Brito et al. 2017; Hua and Minghai 2017) markers in a mark capture-recapture framework to estimate swamp deer populations in this landscape (Andreotti et al. 2016; Kierepka et al. 2016; Sethi et al. 2016; Viengkone et al. 2016; Blåhed et al. 2019; Cook et al. 2020; Li et al. 2020) and assess landscape-scale genetic and demographic parameters, as other standard population estimation approaches such as Camera Trap, Line Transect- (Andriolo et al. 2005; Fragoso et al. 2016; Meek et al. 2019; Paul et al. 2019). are not conducive in the human-dominated landscape.

The extensive swamp deer genetic sampling throughout this landscape has provided some unexpected insights into the species genetic connectivity and health in the Gangetic habitat block. Earlier studies and information suggested that the swamp deer population in this region is small, scattered and possibly inbred (due to limited connectivity through human-dominated areas) (Paul et al. 2018). However, our results clearly disproved such notions regarding swamp deer movement patterns through the geneflow analyses. The weak genetic structure, random spatially-distinct distribution of some highly-related individuals (n=16 individuals (13M: 3F), r=0.6) and no isolation by distance patterns reflect movement events despite fragmentation in this landscape. Although very small numbers of related individuals are found in this study, the implications are very important for this highly-fragmented natural patches of grassland habitats. Large number of studies conducted on various terrestrial mammalian systems report a direct relationship between habitat loss/fragmentation and reduction in genetic connectivity (carnivore- Riley et al. 2006; Carvalho et al. 2018; herbivore- Niedziałkowska et al. 2012; herbivore- (Fraser et al. 2019); omnivore- (Sato et al. 2014)), but several works on ungulates corroborate our findings (Caribou- Boulet et al. 2007; Mager et al. 2013 White-tailed deer- Blanchong et al. 2013; African Buffalo- Epps et al. 2013). Such findings often result from a complex interplay between the environment dynamics and the species ability to disperse and breed successfully (Ito et al. 2013; Mager et al. 2013). The radio-telemetry data confirmed these patterns by providing fine-scale insights on swamp deer dispersal and intensive use of the grassland patches as ‘stopover sites’ during their movement. Net Squared Displacement (NSD) results suggested that the two individuals exhibited migratory (Female 2) and mixed migratory (Female 1) patterns, respectively. The stopover sites identified on the movement routes of both individuals between JJCR and HWLS probably act as important refugia for swamp deer and aid in maintaining genetic connectivity (as evident from the 2^nd^ individual movement data (Fig. 3b)). These results support earlier reports of swamp deer congregations during summer months (to feed on young vegetation in the floodplains) and migrations at the onset of monsoon (Schaaf 1978; Qureshi et al. 2004). Further, the sugarcane fields possibly help in movement between stopover sights in certain time of the year (Paul et al. 2021), as reported from other studies from India (Wikramanayake et al. 2004; Athreya et al. 2007, 2013; Talukdar and Sinha 2013; Warrier et al. 2020). It is however important to realise that these inferences are based on only two collared individual females, and future efforts to radio-tag more animals (including both male and females) from different parts of this landscape could help us to ascertain the main drivers of such seasonal movement events. The spatially exhaustive, homogenous sampling comprising both male (n=219 individuals) and female (n=47 individuals) based analyses also indicate non-biased, gender-common movement pattern, as reported in other cervids (White-tailed deer- Long et al. 2005, 2008, Roe deer- Gaillard et al. 2008; Bonnot et al. 2010, Red deer- Perez-Espona et al. 2010. Another encouraging result from this study that has significant swamp deer conservation implications is the moderate heterozygosity and low inbreeding status of this population. The heterozygosity value ranged between 0.38 to 0.84 across loci (Average Ho= 0.51, SD=0.10) and is consistent with other deer species (Kuehn et al. 2003; Feulner et al. 2004; Lee et al. 2015; Mukesh et al. 2015), including previous studies on swamp deer (Kumar et al. 2017). However, the low inbreeding value (FIS) contradicts earlier reports from JJCR (Kumar et al. 2017) which forms only a part of the entire Ganges population. This pattern of genetic admixture, moderate heterozygosity, relatively low inbreeding and random spatial organisation of related individuals can be explained to some extent by the species biology and history of this landscape. Till the 1940s-1950s, impenetrable swamps and high incidences of malaria infestation made this region inhabitable. The subsequent eradication of malaria with introduction of DDT (as an anti-mosquito agent) and major resettlement of people from erstwhile East Pakistan by the Government of India (Johnsingh et al. 2004) resulted in a sharp human-population increase. Severe encroachment of grasslands for agriculture has led to decline of these swampy grassland habitats from the 70s and 80s (Johnsingh et al. 2004). Despite such loss of habitats, the seasonal movement behaviour of swamp deer (Martin and Gopal 2015) helps maintaining genetic mixing among these fragmented grassland patches, which has earlier been reported as their breeding grounds (Paul et al. 2021). This is also evident from the patterns of very low pairwise FST values (between Zones 1 to 5). Manifestations of genetic discontinuity might take a longer period as it is reported that Fst has a lag time of about 200 generations before it can be detected due to the formation of new barriers (Landguth et al. 2010). IBD tests have a much shorter lag period (1–15 generations) to detect barrier effects (Landguth et al. 2010), but for a vagile species like swamp deer we did not expect to observe impacts of IBD given the movement patterns seen in this relatively small landscape. Our results indicate presence of two intermixing swamp deer genetic lineages along the Gangetic habitat blocks and further efforts are required to understand their origin and status by sampling the Sharda habitat block, which is the largest population of northern swamp deer. Additionally, preliminary evidences also indicate slightly different genetic signatures from the samples collected from the southern part of the study area (zone 5). This area is known to harbour very low-density swamp deer-occurring habitats (Mondol et al. 2019; Paul et al. 2020) and further sampling from these areas is required to ascertain the actual patterns.

## 5. Conservation implications

Our genetic and radio-collaring data suggest that despite severe anthropogenic pressures between JJCR and HWLS, the swamp deer population is connected, retains moderate genetic diversity, and exhibits low levels of inbreeding. These are encouraging signatures for a small, fragmented and isolated cervid population and should be considered carefully for appropriate conservation/management plans. It is important to understand that even though the landscape continues to be functional (active geneflow and animal movement), maintaining the integrity and functionality with very high human density (1164 people/km^2^- (Census of India 2011) and associated anthropogenic activities will remain the most important challenge in future. Recent reports indicate ∼57% loss of grassland habitats (to agricultural conversion) along the upper Gangetic plains during last 30 years (Paul et al. 2021), and therefore conservation of the identified ‘stopover sites’ is absolutely critical as landscape changes can impact gene flow in fragmented landscapes (Fraser et al. 2019). Our field-based mapping and radio-telemetry data points show that majority of the heavily-used stopover sites (at least from the two collared animals) are found in the non-protected areas bordering the states of Uttarakhand and Uttar Pradesh. Therefore, we recommend a jointly-prepared protection and grassland recovery plan by Uttarakhand and Uttar Pradesh Forest departments to control encroachment and grazing pressures on these grassland patches and ensure functional connectivity between JJCR and HWLS. Our intensive survey efforts and subsequent identification of individuals indicate an unequal swamp deer distribution in this landscape, where the area above Bijnor Barrage harbours more individuals (n=216 individuals in 1714 km^2^) compared to below the Barrage (n=50 individuals in 1459 km^2^). This corroborates with the available habitat loss information (∼50% loss in upper part of Bijnor Barrage and ∼65% loss below Barrage area) (Paul et al. 2021). Recent conservation initiatives have identified “Priority Conservation Areas” (Paul et al. 2020) in this landscape and ensured government approval of HWLS boundary reappropriation (Mondol et al. 2019) and therefore it is critical to focus on the management of the lower part of Bijnor Barrage. The Gangetic ecosystem is highly dynamic and protecting the grasslands would require collaborative efforts involving multiple stakeholders including several government departments (Ministry of Agriculture and Farmers Welfare, Ministry of Housing and Urban Affairs, Department of Water Resources, River Development and Ganga Rejuvenation, Department of Revenue etc.) to strategize appropriate habitat recovery, plantation management, distribution of minimum numbers of agricultural licenses along rivers, review of the land tenure/revenue records, the release of water from dams/barrages etc. Such comprehensive effort only can ensure long-term viability of these productive habitats.

Around 5,000 swamp deer remain in the wild globally (Duckworth et al. 2015), but currently the focus of their conservation is limited to the protected areas (Mondol et al. 2019; Paul et al. 2020). The Gangetic habitat population represents the western-most distribution of the species and the future is promising, provided that connectivity is maintained and habitat management becomes an important conservation agenda in immediate future. We hope that the results presented in this study would bring out the conservation attention to all concerned stakeholders and help ensuring the long-term persistence of this species outside protected area habitats. Given that a large number of species are distributed outside protected areas globally, this study could become an example to deal with the conservation challenges faced by them.

## Statements and Declarations

### Author contributions

SM, DM, BP conceived the study idea. SM and BP generated funds and supervised the study. BH procured the collars. PN spearheaded the collaring-operation. SP, BP, SM, DM and BH all supported in collaring-operation. SP conducted sampling and data generation. SS and GP supported in data generation and analysis. SP and SM wrote the initial manuscript. SP, SM and SS revised the draft and all authors approved the final draft.

### Data availability

The data generated in this study is available in Figshare (https://figshare.com/s/fe0cb79541ca8b3ae2ea)

### Competing interests

The authors have not disclosed any competing interests

#### Acknowledgement

We acknowledge the Forest Departments of Uttarakhand, Uttar Pradesh and Ministry of Environment, Forest and Climate Change (No. 244/2018/RE), Government of India for providing necessary permits and support to carry out our field work. Our thanks to the Forest Department officials especially Divisional Forest Officer, Haridwar Forest Division and local community members for assisting in sample collection. We acknowledge help from Vinod Thakur, Tista Ghosh, Suvankar Biswas, Dr. Sarbesh Rai, Shiv Kumari Patel, Shrushti Modi, Kaushal Singh Sissodia, Sultan Singh Badana, Nimisha Srivastava, Rakesh Mondol, Debanjan Sarkar, Prajak Kumar Das, Chitrapal, Imam, Ranju, Bhura, Annu, Juri, Inam, Ammi, M.Sc. students of Wildlife Institute of India (XVI batch) and villagers of Gendikhata and Tantwala for their help during the collaring operation and field surveys. We acknowledge Prof Paulo Alves and Dr Joao Queiros from University of Porto, Portugal for providing the STR markers. We appreciate technical help in laboratory from Tista Ghosh, Mouli Bose, Aamer Sohel Khan and A. Madhanraj. We thank the Director, Dean, Research Coordinator and Nodal Officer of Wildlife Forensics and Conservation Genetics cell of Wildlife Institute of India for their support. The work was funded by Uttarakhand Forest Department, Uttar Pradesh Forest Department, Ministry of Environment, Forest and Climate Change, Government of India and Conservation Force. Shrutarshi Paul was awarded Department of Science and Technology INSPIRE Research Fellowship (IF150680) and Samrat Mondol was supported by Department of Science and Technology INSPIRE Faculty Award (IFA12-LSBM-47).

---

**Supplementary Table 1:** Details (Grasslands, Blocks, Districts, States) of all locations of the antlers and pellets collected in this study.

| Sl No. | Name of the area | Block | District, State | No. of antlers | No. of pellets |
| --- | --- | --- | --- | --- | --- |
| 1 | Jhilmil Jheel Conservation Reserve (JJCR) | Block 1 | Haridwar, UK | 88 | 41 |
| 2 | Bhogpur | Block 1 | Haridwar, UK | 0 | 0 |
| 3 | Banganga khadar | Block 1 | Haridwar, UK | 0 | 0 |
| 4 | Ranjeetpur and adjacent river islands | Block 1 | Haridwar, UK & Bijnor, UP | 4 | 17 |
| 5 | Amichand and Nangal | Block 1 | Haridwar, UK & Bijnor, UP | 4 | 15 |
| 6 | Dharampur- Jogga Jheel Complex | Block 1 & Block 2 | Haridwar, UK & Muzaffarnagar, UP | 15 | 9 |
| 7 | Sukhapur- Manwala- Chandpuri complex | Block 2 | Bijnor, UP | 0 | 21 |
| 8 | Mirapur khadar | Block 2 | Bijnor, UP | 0 | 0 |
| 9 | Shukratal- Haiderpur complex | Block 2 & Block 3 | Muzaffarnagar & Bijnor, UP | 141 | 36 |
| 10 | Jeevanpuri and adjacent river islands | Block 3 | Bijnor, UP | 1 | 13 |
| 11 | Buriganga jheel | Block 3 | Meerut, UP | 0 | 2 |
| 12 | Dharmapura- Daranagarganj complex | Block 3 | Muzaffarnagar & Bijnor, UP | 1 | 16 |
| 13 | Jahanabad- Joagli jheel complex | Block 3 & Block 4 | Bijnor, UP | 0 | 6 |
| 14 | Hadipur Gori | Block 4 | Meerut, UP | 0 | 9 |
| 15 | Rahmanpur- Jahangirpur complex | Block 4 | Bijnor, UP | 0 | 2 |
| 16 | Jalalpur khadar | Block 4 | Meerut, UP | 1 | 27 |
| 17 | Kalidab khadar | Block 4 & Block 5 | Amroha, UP | 0 | 0 |
| 18 | Kopla jheel | Block 5 | Hapur, UP | 3 | 16 |

\*UK= Uttarakhand; UP= Uttar Pradesh

**Supplementary Table 2:** Details of the initial set of 48 primers tested for swamp deer individual identification.

| SI No. | Marker Name | Selection Status | Reason for Rejection |
| --- | --- | --- | --- |
| 1 | CEH 4 | Not selected | Multiple bands |
| 2 | CEH 22 | Not selected | Multiple bands |
| 3 | CEH 45 | Not selected | Multiple bands |
| 4 | CEH 60 | Not selected | Multiple bands |
| 5 | CEH 73 | Not selected | Multiple bands |
| 6 | CEH 79 | Not selected | Multiple bands |
| 7 | CEH 63 | Not selected | Multiple bands |
| 8 | CEH 15 | Not selected | Multiple bands |
| 9 | CEH 44 | Not selected | Multiple bands |
| 10 | CEH 80 | Not selected | Multiple bands |
| 11 | CEH 81 | Not selected | Multiple bands |
| 12 | CEH 6 | Not selected | Multiple bands |
| 13 | CEH 38 | Not selected | Multiple bands |
| 14 | CEH 78 | Not selected | Multiple bands |
| 15 | CEH 39 | Not selected | Multiple bands |
| 16 | CEH 23 | Not selected | Multiple bands |
| 17 | CEH 57 | Not selected | No amplification |
| 18 | CEH 85 | Not selected | No amplification |
| 19 | CEH 31 | Not selected | Non-specific bands |
| 20 | CEH 55 | Not selected | Non-specific bands |
| 21 | CEH 72 | Not selected | Non-specific bands |
| 22 | CEH 70 | Not selected | Non-specific bands |
| 23 | CEH 28 | Not selected | Inconsistent amplification |
| 24 | CEH 49 | Not selected | Inconsistent amplification |
| 25 | CEH 86 | Not selected | Inconsistent amplification |
| 26 | CEH 34 | Not selected | Inconsistent amplification |
| 27 | CEH 48 | Not selected | Inconsistent amplification |
| 28 | CEH 68 | Not selected | Excessive stutter bands and low RFUs |
| 29 | CEH 74 | Not selected | Excessive stutter bands and low RFUs |
| 30 | CEH 87 | Not selected | Unstable allele characteristics |
| 31 | CEH 77 | Not selected | Unstable allele characteristics |
| 32 | CEH 84 | Not selected | Unstable allele characteristics |
| 33 | CEH 61 | Not selected | Inconsistent results with antler and pellet DNA |
| 34 | CEH 64 | Not selected | Inconsistent results with antler and pellet DNA |
| 35 | CEH 76 | Not selected | Inconsistent results with antler and pellet DNA |
| 36 | CEH 53 | Selected | - |
| 37 | CEH 58 | Selected | - |
| 38 | CEH 71 | Selected | - |
| 39 | CEH 35 | Selected | - |
| 40 | CEH 33 | Selected | - |
| 41 | CEH 43 | Selected | - |
| 42 | CEH 52 | Selected | - |
| 43 | CEH 50 | Selected | - |
| 44 | CEH 82 | Selected | - |
| 45 | CEH 56 | Selected | - |
| 46 | CEH 51 | Selected | - |
| 47 | CEH 47 | Selected | - |
| 48 | CEH 75 | Selected | - |

